# Genome-wide identification and transcriptomic analysis of microRNAs across various amphioxus organs using deep sequencing

**DOI:** 10.1101/736371

**Authors:** Qi-Lin Zhang, Hong Wang, Qian-Hua Zhu, Xiao-Xue Wang, Yi-Min Li, Jun-Yuan Chen, Hideaki Morikawa, Lin-Feng Yang, Yu-Jun Wang

**Author notes:** Correspondence (Y.J.W) or (Q.L.Z) or (L.F.Y). These authors contributed equally (Qi-Lin Zhang, Hong Wang, Qian-Hua Zhu, Xiao-Xue Wang).

## Abstract

Amphioxus is the closest living invertebrate proxy of the vertebrate ancestor. Systematic gene identification and expression profile analysis of amphioxus organs is thus important for clarifying the molecular mechanisms of organ function formation and further understanding the evolutionary origin of organs and genes in vertebrates. The precise regulation of microRNAs (miRNAs) is crucial for the functional specification and differentiation of organs. In particular, those miRNAs that are expressed specifically in organs (OSMs) play key roles in organ identity, differentiation, and function. In this study, the genome-wide miRNA transcriptome was analyzed in eight organs of adult amphioxus *Branchiostoma belcheri* using deep sequencing. A total of 167 known miRNAs and 23 novel miRNAs (named novel_mir), including 139 conserved miRNAs, were discovered, and 79 of these were identified as OSMs. Additionally, analyses of the expression patterns of eight randomly selected known miRNAs demonstrated the accuracy of the miRNA deep sequencing that was used in this study. Furthermore, potentially OSM-regulated genes were predicted for each organ type. Functional enrichment of these predicted targets, as well as further functional analyses of known OSMs, was conducted. We found that the OSMs were potentially to be involved in organ specific functions, such as epidermis development, gonad development, muscle cell development, proteolysis, lipid metabolism and generation of neurons. Moreover, OSMs with non-organ specific functions were detected, and primarily include those related to innate immunity and response to stimuli. These findings provide insights into the regulatory roles of OSMs in various amphioxus organs.

## INTRODUCTION

The phylum Chordata contains three subphyla members, including the Cephalochordata (e.g. amphioxus), Urochorda (e.g. ciona), and Vertebrata (Satoh *et al*., 2014). The origin and evolution of organs and tissues in chordates is one of the most important scientific questions in evolutionary developmental biology (evo-devo) (Holland and Chen, 2001). However, with the exception of a recent report by Marlétaz *et al*. (2018),, genetic information from different organs of amphioxus as the sister lineage to all other chordates remain largely unstudied, which limits our understanding for molecular mechanisms of organ function formation in chordate as a vertebrate outgroup and origin of organ complexity in vertebrates. Amphioxus, also known as lancelets, which belong to the subphylum Cephalochordate, retains some of the structural morphology characteristics observed in common ancestors between cephalochordates and vertebrates from the Cambrian period of ∼530 million years ago (Chen *et al*., 1999; Putnam *et al*., 2008). Therefore, cephalochordates are key experimental animals for the study of evo-devo, evolutionary origins of organs, and comparative immunology of vertebrates (Putnam *et al*., 2008; Huang *et al*., 2014).

Non-coding RNAs (ncRNAs) regulate many biological processes, such as development, apoptosis, metabolism, differentiation, and the function formation of cells in animals (Arner and Kulyté, 2015). Of the ncRNAs, those known as microRNA (miRNA, ∼22 nucleotides) are the most extensively investigated types. In general, many of these miRNAs are transcribed from introns of protein-coding genes, and negatively regulate expression of their targets by complementary base-pairing to regions in the 3’ UTR (untranslated regions) of mRNAs in animals (Campo-Paysaa *et al*., 2011). Furthermore, miRNAs may contribute to the function formation of different organs through their specific expression in amphioxus. It has been reported that miR-92 is specifically expressed in the hepatic caecum, gill, and intestines in amphioxus. It is believed to regulate the complement pathway by targeting C3 to promote the immune response against bacterial infection (Yang *et al*., 2013). Muscle-specific expression of miR-1 and miR-133 was clearly demonstrated using *in situ* hybridizationusing in amphioxus early larvae (Campo-Paysaa *et al*., 2011). Candiani *et al*. (2011) identified six miRNAs with specific expression in the nervous system of amphioxus by whole-mount *in situ* hybridization. Both differentiating and mature neurons exhibited specific expression of miR-124. Thus, the body of literature suggests that miRNAs may be key regulatory molecules for the regional and specific function formation of different organs in amphioxus. So far, only miRNA expression profiles in three digestive and immune-related organs, gill, intestine and hepatic caecum, have been obtained using microarray technology based on nucleic acid hybridization (Liao *et al*., 2017a). However, only three organ types were used and microarray based on non-sequencing technology is not feasible for the identification of novel molecules in different organs (Gao *et al*., 2011), possibly hindering a systematic and accurate discovery and function analysis of organ-specific expressed genes.

Next-generation sequencing is a useful tool for the analysis of the expression profile of miRNAs. Its advantages are unbiased large-scale detection of small RNAs at a genome-wide level, even for transcripts with low expression levels, and its ability to identify novel RNA miRNAs (Yao *et al*., 2012). To date, there has not been a comparative analysis of miRNA expression profiles among different organs in amphioxus using deep sequencing. In this study, deep sequencing was used to systematically analyze miRNAs purified from eight organs of adult Chinese amphioxus (*Branchiostoma belcheri*). In addition to demonstrate organ-specific expression of a large group of known miRNAs and novel miRNA candidates among different organs, the corresponding target genes of these miRNAs were predicted, and the function of known organ-specific expressed miRNAs was explored. Then, the function of all organ-specific expressed miRNAs (OSMs) was analyzed by enrichment of Gene Ontology (GO) terms using bioinformatics. The dataset obtained here will be a valuable resource for uncovering the potential roles of miRNAs in organ differentiation of amphioxus.

## MATERIALS AND METHODS

### Ethics statement

This study was carried out in accordance with the recommendations of the Guide for the Care and Use of Laboratory Invertebrate Animals. The protocol was approved by the Ethical Committee of Researches of the Nanjing University.

### Sample preparation

Healthy adult individuals of *B. belcheri* were obtained from the Evo-devo Institute of Nanjing University at Beihai City, Guangxi Province, China. Experimental animals were kept in acrylic tanks with filtered seawater, following our previously used methods (Liao *et al*., 2017b). These individuals were maintained for several days to empty their intestinal and hepatic caecum contents. Subsequently, each of the approximately 40 individuals was placed on ice and dissected to collect the eight specific organs (i.e. nerve cord, notochord, skin, intestine, hepatic caecum, muscle, gill and ovary) used in this study. Each organ type obtained from different individuals was pooled together in a 1.5 ml RNase-free microcentrifuge tube, and immediately frozen using liquid nitrogen and stored at −80 °C. All experimental samples were stored at Beihai Marine Station of Nanjing University in Beihai, Guangxi province, China.

Paraffin section of *B. belcheri* was generated according to previous methods (Yang *et al*., 2013). In brief, adult *B. belcheri* was fixed in 4% paraformaldehyde at 4 °C for 12 hours, dehydrated with graduated ethanol, and embedded in paraffin. Then, the embedded amphioxus was cut into 10 μm sections. After deparaffinization, organ sections were dyed using Hematoxylin and Eosin Staining Kit (Yeasen, USA) following the manufacture’s manual. Next, the dyed sections were photographed by Olympus microscope DP71 (Olympus, Japan).

### RNA extraction, library construction, and sequencing

The total RNA of each organs type was extracted using Trizol reagent (Invitrogen, USA), according to the manufacturer’s protocol. Residual DNA contamination was removed by RNase-free DNase (Qiagen, Germany). RNA concentration and quality were initially assessed using a BioPhotometer Plus (Eppendorf, Germany). Next, an Agilent 2100 Bioanalyzer (Agilent Technologies, USA) was employed to further verify RNA structural integrity and quality. RNA samples with a RNA integrity number (RIN value) ≥ 7 were used in further experiments. Preparation of small-RNA libraries was performed using Illumina TruSeq Small RNA Library Preparation Kits (Illumina, USA). Quality of the small-RNA library was assessed using the Agilent 2100 Bioanalyzer (Agilent Technologies, USA) and an ABI StepOnePlus Real-Time PCR System (Applied Biosystems, USA). Deep sequencing of the eight prepared libraries was performed on an Illumina HiSeq4000 platform (Illumina, USA) with 50-bp single-end reads at the Beijing Genomics Institute (BGI-Shenzhen, China).

### Analysis of deep sequencing data

Clean reads were obtained from the raw data by removing low-quality tags, reads with poly-N tails or 5′ adapter contaminants, reads with 3′ null adapters, and insert null tags. Next, clean small RNA (sRNA) tags were mapped to the *B. belcheri* genome (v18h27.r3; http://genome.bucm.edu.cn/lancelet/, early available) using Bowtie2 (parameter: -q -L 16 --phred64 -p 6) to analyze their expression and distribution across the reference genome (Langmead *et al*., 2009). To discard tags originating from rRNAs, tRNAs, snRNAs, snoRNAs, and sRNA tags, a BLAST search was conducted against the Rfam 11.0 and the NCBI GenBank databases. RepeatMasker software and genome-based mapping information were used to remove tags from repeat regions and protein coding sequences (Zhong *et al*., 2015). The sRNA sequences were identified by performing a BLAST search in miRBase21.0 (Griffiths-Jones, 2006), allowing a maximum of two mismatches to known miRNA.

### Identification of novel miRNAs and homologous miRNAs

The remaining sRNA tags originating from exon sense and intron and intergenic regions were used to identify novel miRNAs and their precursors. The prediction software mirDeep2 was used to identify novel miRNAs using default options recommended by software manual (Friedländer *et al*., 2012). MirDeep2 has a substantially improved algorithm for the identification of miRNAs from RNA sequencing data of animal groups and can predict canonical and non-canonical miRNAs with an accuracy of 98.6-99.9% (Friedländer *et al*., 2008; Friedländer *et al*., 2012). To ensure the accurate discovery of novel miRNAs through verification of the secondary structures, novel miRNA precursors (pre-miRNAs) were analyzed using RNAfold (Höner et al., 2011) to estimate whether the secondary structure of the precursor is a perfect stem-loop formation. Furthermore, the predicted novel miRNA tags/sequences with mapped tags of less than 5 were discarded (Zhang *et al*., 2017a), and the remainder retained as the novel mirDeep2 miRNAs. Meanwhile, the additional prediction software Mireap (Yuan *et al*., 2013) was used to discover novel miRNAs by predicting the hairpin structure, dicer cleavage site and minimum free energy of the unannotated small RNA tags that had been mapped to the genome. Those tags with a minimum free energy (MFE) value of the folding precursor ≤ −19 kcal/mol were kept as novel miRNAs identified by Mireap. The intersection of identified miRNAs by mirDeep2 and Mireap was retained. Then, these identified miRNAs were further filtered as the final novel miRNA set under strict criteria (Huang *et al*., 2017): 1) the presence of both 5p and 3p strands in the read dataset; 2) their overlap with a ∼2 nt overhang on each side when the hairpin is folded. Subsequently, novel miRNAs and *B. belcheri* known miRNAs above annotated were searched in miRBase21.0 animal database (excluding *B. belcheri* dataset), allowing for a maximum of two mismatches and those that were annotated as known miRNAs of other animals were considered to be conserved/homologous.

### Normalization of miRNAs expression and identification of organ-specific miRNAs

Expression levels of all miRNAs (conserved and novel miRNAs) were normalized using TPM values (Zhong *et al*., 2015). To estimate the organ specificity of a miRNA, an entropy-based metric that relies on Jensen-Shannon (JS) divergence was employed to calculate specificity scores (ranging from 0 to 1) based on previous descriptions of the JS algorithm (Cabili *et al*., 2011). This specificity metric quantifies the similarity between a transcript’s expression pattern across tissues and another predefined pattern that represents the extreme case in which a transcript is expressed only in one tissue. Therefore, a perfect organ-specific expression pattern is scored as JS = 1, indicating a transcript is expressed only in one organ type (Cabili *et al*., 2011). Specifically, index τ was calculated according to previously published methods using a custom Perl script to detect the organ specificity of miRNA expression (Yanai *et al*., 2005). For a given transcript, the index τ is defined as 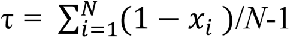, where *N* is the number of *x*_*i*_ organs, and *i*^*th*^ is the expression value in the *i*^th^ organ normalized by the maximal expression value across all of the organs (Li *et al*., 2015). First, it was determined whether expression was organ-specific for each miRNA. If miRNAs were observed to have values for τ > 0.85, they were considered to be expressed in an organ-specific manner (organ-specific miRNAs, OSMs) (Cabili *et al*., 2011). Next, if OSMs were observed, JS scores were used to evaluate organs for the specific expression for each OSM as previously described (Cabili *et al*., 2011; Li *et al*., 2015). Finally, each τ value was compared with the other JS scores (corresponding to each of the eight organs) for each miRNA. Organ specificity of miRNA expression and the respective expression-specific organs were successively confirmed.

### Determination and functional analysis of potential miRNA targets

The potential target genes of OSMs were predicted using two bioinformatic tools. The first was RNAhybrid (Kruger et and Rehmsmeier, 2006), and the default parameters were used as follows: -b 100 -c -f 2,8 -m 100000 -v 3 -u 3 -e -20 -p 1 -s 3utr_Bb. The second tool used was miRanda (Betel et al., 2008), with the following default parameters of -en -20 –strict being used. The intersection point for the results from two prediction tools was assumed to be the set of reliably predicted target genes of the OSMs. These potential targets were selected for downstream analyses. To determine the functions of the predicted OSM regulated target genes, the genes were predicted target genes of all miRNAs identified in this study were annotated using a basic local BlastX tool (Kent, 2002), using the default parameters in the NCBI non-redundant (NR) animal database. To further understand the regulatory function of OSMs in different organs of *B. belcheri*, known OSM (analyzed based on *B. belcheri* miRNAs previously identified and stored at miRBase database) target genes predicted by bioinformatics were filtered and extracted according to the following criteria: 1) target genes with unknown annotation and hypothetical proteins were removed; and 2) mutiple targets with the same gene description were filtered, with only one being retained as the representative. Next, GO annotation of each gene was extracted using Blast2GO pipeline (Conesa *et al*., 2005) according to the NR annotation. Because annotation of model animals is full, while that of non-model species is lack, thus sequences of *B. belcheri* were searched in libraries of all the animal species in the GO term analysis to obtain adequately best match. We performed the GO enrichment analysis for the predicted target genes of the OSMs using Fisher’s exact test in Blast2GO pipeline (Conesa *et al*., 2005), and outputs (*P*-values) of the software were used to perform multiple test corrections by false discovery rate (FDR). GO terms presenting FDR< 0.05 were retained. The redundant GO terms were then reduced using the GO trimming tool (Koop et al., 2011). GO terms for biological processes were retained to represent the enriched functions of predicted target gene clusters.

### Quantitative real-time PCR

Collection of the eight organs and extraction of their total RNA, including small RNAs, were performed according to the above-mentioned methods. For each organ type, three biological replications were used. Mature miRNA expression was measured using Taqman probe kits (Applied Biosystems, USA) customized for each gene, including the RT primers, PCR primers and Taqman probes. The product information for each miRNA probe is presented in Supplementary File 1. We selected U6 as a housekeeping gene. Each reaction was conducted in three technical replicates. Relative expression levels of target miRNAs were normalized using the 2^−ΔΔCt^ method (Livak and Schmittgen, 2001). The data shown are presented as the mean ± SD and figures were generated by SigmaPlot 12.0. According to methods described in our previous studies (Zhang *et al*., 2017b), consistency of miRNA expression patterns between deep sequencing and qRT-PCR were evaluated by calculating the Pearson’s correlation coefficient and its significance (*p*-value) level in IBM SPSS statistics 22 software.

## RESULTS

### Overall assessment of the miRNAome of different organs

Approximately 34 million clean reads for each of the eight organ samples, including skin, ovary, notochord, nerve cord, muscle, intestine, hepatic caecum, and gill, were obtained after filtering low-quality reads and contaminants (Figure 1A, B, C and Supplementary File 2). All of the organ samples showed peak length of the miRNAs at about 22-23nt (Supplementary File 3), as has been previously described for *B. belcheri* and other animals (Zhang *et al*., 2017a). All sequencing data have been released to the NCBI SRA database under accession number SRR9951226-SRR9951233. The ratios of total clean tags that mapped to the *B. belcheri* genome ranged from 74.61% to 82.23%, indicating that a set of reliable clean tags was obtained (Zhang *et al*., 2017a). To minimize false positive signals, only tags with Transcripts Per Million (TPM) values ≥ 5 were retained for further bioinformatic analysis. Based on the annotated results of sRNA, it is found that miRNAs occupied a high proportion of sRNA tags in *B. belcheri* organs investigated in this study (> 50%, Supplementary File 4), indicating that miRNA molecules were adequately enriched.

**Figure 1.**
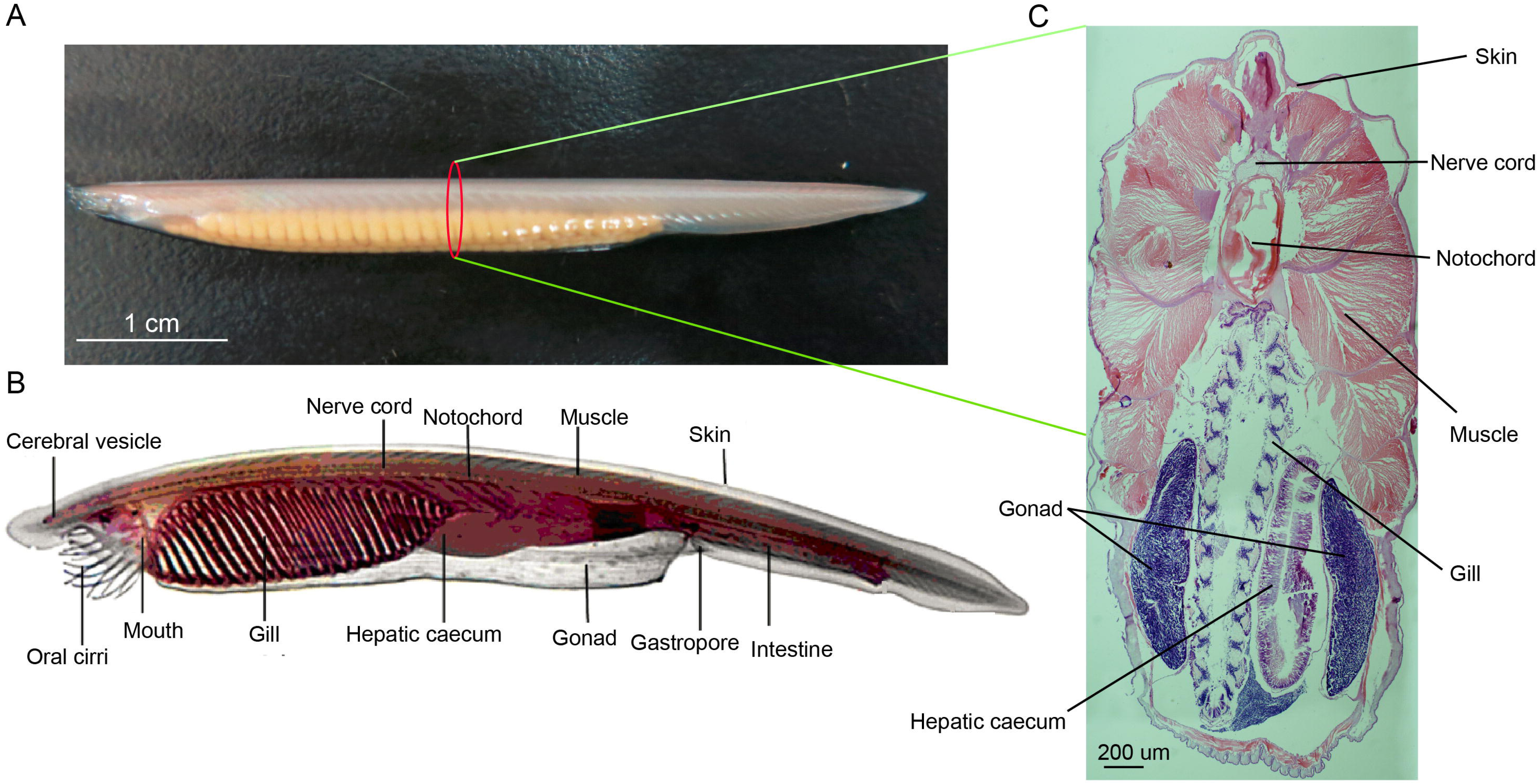
**(A)** Female adult amphioxus *Branchiostoma belcheri* (photographed by EOS 6D digital single-lens reflex cameras (Canon, Japan)). **(B)** An overview of body structure of adult *B. belcheri* (adapted from http://faculty.baruch.cuny.edu/jwahlert/bio1003/chordata.html). **(C)** Hematoxylin and eosin (H&E) staining on transverse sections of an adult *Branchiostoma belcheri*, with staining of nucleus purple and the cytoplasm pink.

### Identification of known, conserved and potential novel miRNAs in *B. belcheri*

A total of 167 known miRNAs, distributed amongst 82 miRNA families, were expressed in at least one organ type (Table 1). In miRBase21.0, 173 mature bbe-miRNA sequences have been stored, and approximately 97% of these were identified in the current study. Of these previously annotated miRNAs, approximately 93% are expressed in the nerve cord. Moreover, 138 known miRNAs were identified to be homologous with those of *B. floridae*, 39 of which are homologous to microRNAs in vertebrates (Supplementary File 5). Together, this indicates that these 39 vertebrate miRNAs have evolved in ancient chordates. However, and 99 of the known miRNAs have only been detected in amphioxus.

**Table 1.** Statistics summary of miRNA in eight organs of *Branchiostoma belcheri*.

| Organ types | Known miRNAs | Novel miRNAs |
| --- | --- | --- |
| Nerve cord | 160 | 10 |
| Notochord | 147 | 8 |
| Skin | 154 | 8 |
| Intestine | 151 | 8 |
| Hepatic caecum | 152 | 12 |
| Muscle | 157 | 12 |
| Gill | 155 | 13 |
| Ovary | 138 | 7 |
| Total amount | 167 | 23 |

A total of 23 potential novel miRNAs were shared (expressed in all eight organs) (Supplementary File 6) across all organs sequenced in this study. These were analyzed further for the detection of the organ expression specificity and specifically expressed organs by caculating their respective τ and then JS scores, respectively. Predicted precursor structure of these novel miRNAs is presented in (Supplementary File 7). Furthermore, novel_mir1 was detected, and is conserved with bfl-miR-4871-3p of *B. floridae* (Jin *et al*., 2017; Zhang *et al*., 2017a). Homologs of the remaining novel miRNAs were not detected, indicating that these miRNAs may potentially be lineage-specific in *B. belcheri*. Alternatively, these remaining novel miRNAs may be existed in other species but haven’t been detected yet.

### Identification and functional enrichment analysis of OSMs

Of all *B. belcheri* miRNAs identified here, 79 OSMs (41.58%) showed organ-specific expression (τ > 0.85) (Supplementary File 8). Furthermore, the distribution tendency of maximal JS scores is presented in Figure 2. Among them, 68 OSMs were observed to be conserved with those of *B. floridae* as well as other vertebrates, accounting for 86.08% of all OSMs (Supplementary File 8). Among the 8 organs, the nerve cord sample showed the highest number of OSMs (26), followed by the gill (20), the hepatic caecum (11), the muscle (8), and the skin (7). The ovary (3), the notochord (2) and the intestine (2) had the lowest numbers of OSMs (Figure 3A). All OSMs are listed in Supplementary File 8, including known and novel OSMs. The known OSMs are also presented in Figure 3B.

**Figure 2.**
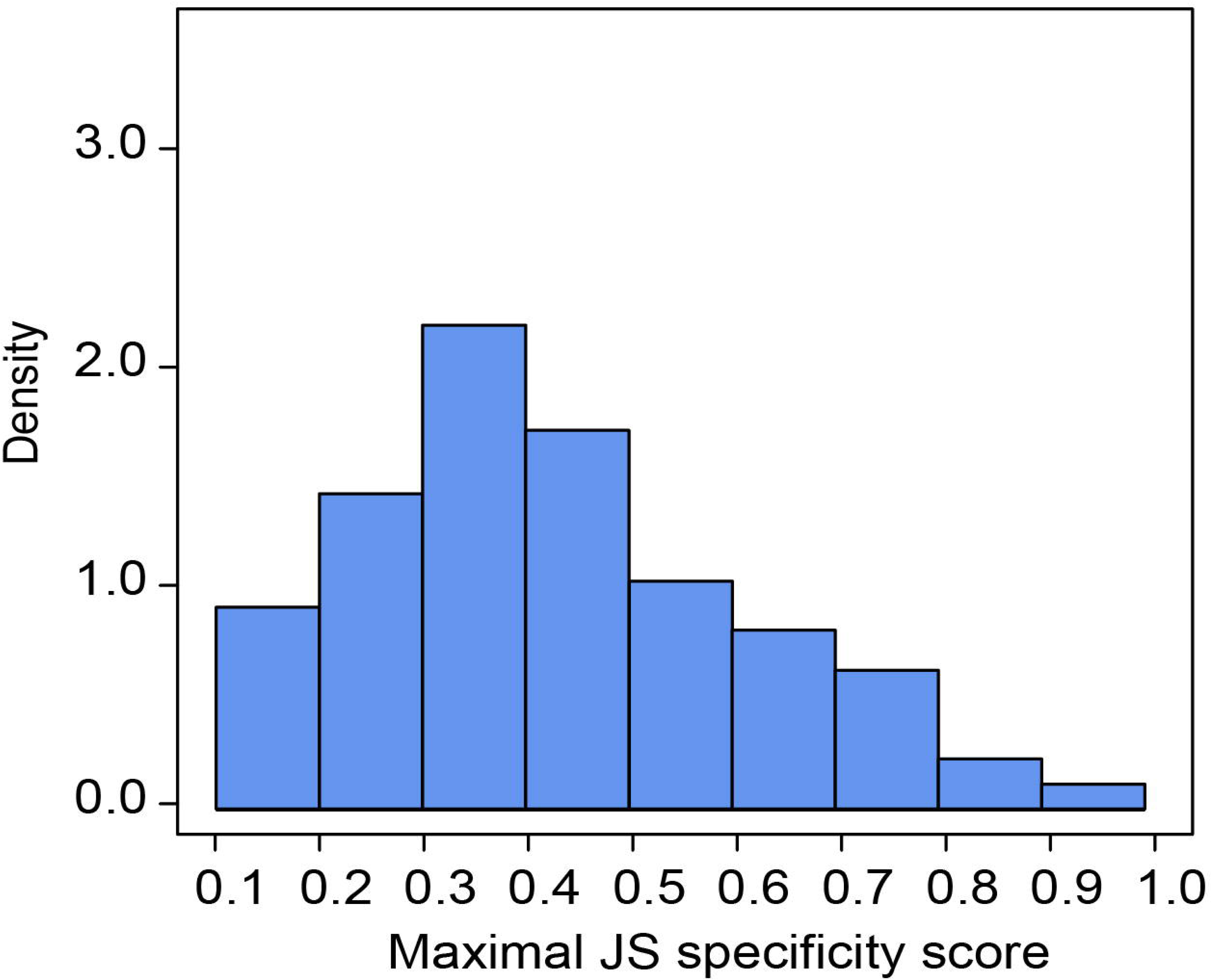
Organ specificity of miRNAs. Distribution of maximal Jensen-Shannon (JS) specificity scores calculated for each organ-specific miRNA (τ > 0.85) across all eight organs.

**Figure 3.**
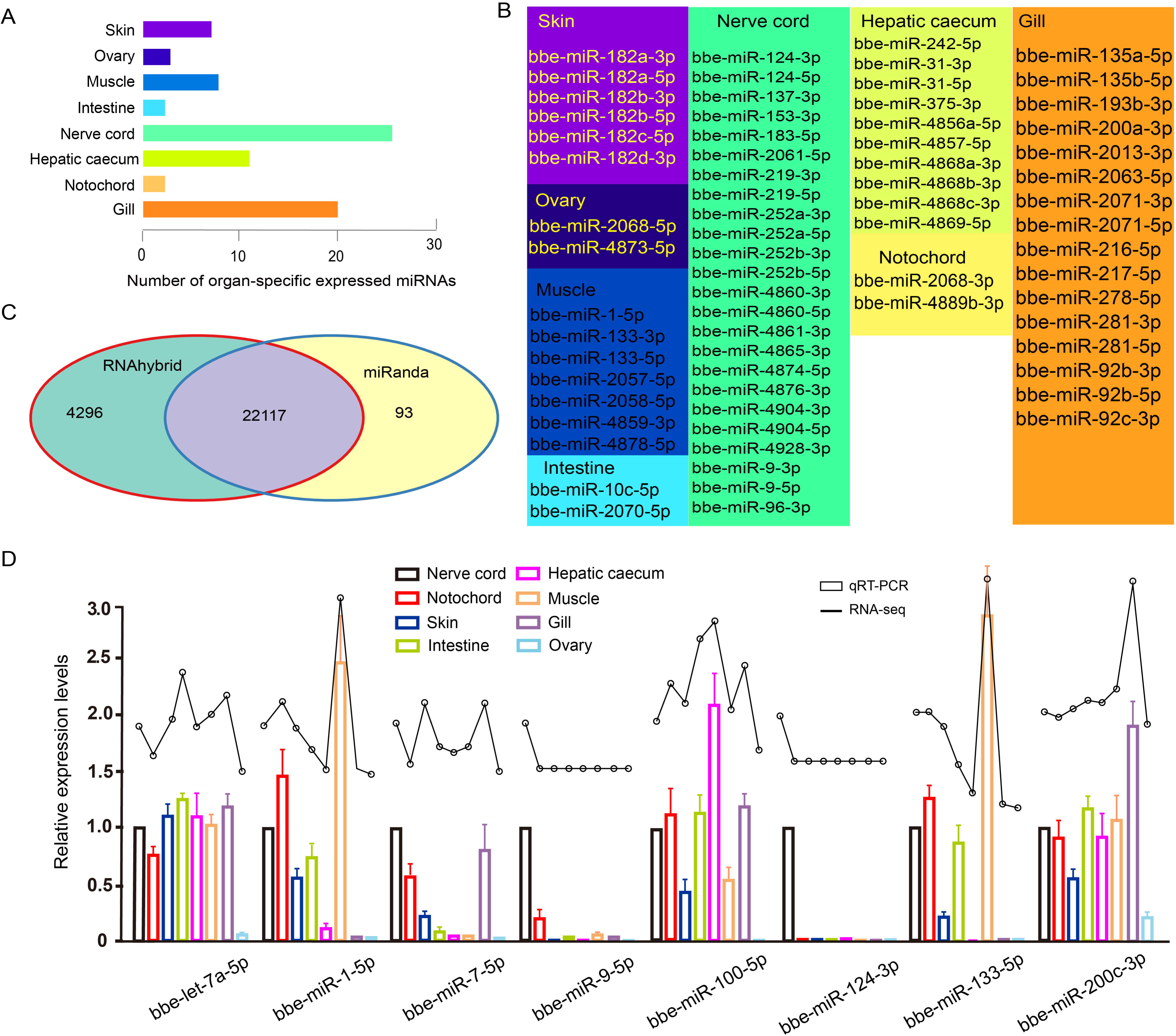
**(A)** Number distribution of organ-specific miRNAs across eight organs. **(B)** The list of known organ-specific miRNAs in eight organs. **(C)** Number of predicted target genes of organ-specific miRNA using RNAhybrid and miRanda software. **(D)** Validation of expression of miRNAs by qRT-PCR experiment. Expression of eight selected randomly miRNAs was detected by qRT-PCR in *B. belcheri* organs including the skin, ovary, muscle, intestine, hepatic caecum, gill, nerve cord, and notochord.

The comprehensive miRNA catalog allows us to explore the potential functions of the miRNAs observed in *B. belchrei*. Here, 22,117 potential targeted genes were identified using both the RNAhybrid and miRanda software programs (Figure 3C). Using GO enrichment analysis, the putative target genes of the OSMs in each organ were to be enriched using the GO terms belonging to the “Biological Process” subcategory (Supplementary File 9). In the skin, 12 enriched GO terms were primarily related to response to stimulus and epidermis development. These terms also included “apoptotic process”, “response to UV-B”, “response to mechanical stimulus”, and “epidermis development”. For the ovary, the 5 GO terms obtained were overrepresented and associated with signaling processes, immune response and ovary development, which included “signaling”, “defense response”, and “gonad development. Unsurprisingly, GO terms enriched from predicted target genes of muscle OSMs were primarily involved in muscle development and morphogenesis. The intestine, hepatic caecum, and gill are considered to be digestive and immune tissues of amphioxus (Han *et al*., 2010; Liu *et al*., 2009). In these three organs, many GO terms involving immune and stimulus response, apoptosis, immune system, metabolic and catabolic processes were widely enriched. Notably, the GO term “proteolysis” was specifically exhibited in the intestine. Meanwhile, it was observed that OSMs of the hepatic caecum also specifically regulated target genes related to stimulus and organic metabolism. In the nerve cord, OSMs were primarily associated with stimulus and drug response, circadian rhythm, sensory behavior, and neural development and function. The GO terms enriched by predicted targets of notochord OSMs were related to cell process and function, phosphorylation and development process.

### Analysis of predicted target genes of known OSMs

Known OSM target genes predicted by bioinformatics were listed (Supplementary File 10). We found that some typical genes involved in stimulus response could be regulated by specific expression of several known OSMs in the skin, such as heat shock protein 70 (*HSP70*) and genes encoding nod-like receptors (NLR protein), mitogen-activated protein kinase 3 (*MAP4K3*), mucin-2-like (*MUC2L*) and tumor necrosis factor receptor superfamily member 11A (*TNFRSF11A*), toll-like receptor 4 (*TLR4*) and collagen alpha-1 (XII) chain (*COL12A1*). Additionally, genes related to calmodulin regulation were found as predicted targets of skin OSMs, including calmodulin-related protein 3 (*CALREP3*), calcium-activated potassium channel subunit alpha-1 (*KCNMA1*), and calcium/calmodulin-dependent protein kinase type II subunit gamma (*CAMK2G*).

In the ovary, bbe-miR-2068-5p was predicted to regulate lipid metabolism function, reactive oxygen species and gene encoding piwi-like 1 protein (*PIWIL1*), while bbe-miR-4873-5p participated probably in the regulation of genes involving stimulus and immune responses, such as mitogen-activated protein kinase kinase kinase15 (*MAP3K15*), WD repeat-containing protein 26 (*WDR26*), and ubiquitin-like modifier-activating enzyme 6 (*UBA*6).

In the notochord, it is found that genes potentially related to notochord development were regulated by known OSMs, including insulin-like growth factors 1 (*IGF1*), bone morphogenetic protein 3 (*BMP3*), notochord homeobox-like protein, muscle, and skeletal receptor tyrosine-protein kinase (*MUSK*).

In the nerve cord, various innate immune-related genes were found to be putative targets of OSMs, including complement (*C1q1*, complement factor H (*CFH*), complement component factor B/C2 (*B/C2*)), pattern recognition receptors (PRRs) (NLR protein, RIG-I-like receptor LGP2), cytokines (interferon regulatory factor 2, 4, 5 (*IRF2*, *4*,*5*)), adaptors and signal transducers (TNF receptor-associated factor 6 (*TRAF6*), and WD repeat-containing protein 6-like protein). Some typical genes associated with nerve development and function were also found, such as fibroblast growth factor receptor (*FGFR2*), neurogenic locus protein delta-like protein, neurogenic locus notch-like protein 3 (*NOTCH3*), paired box protein Pax-6 (*PAX6*), *AMPHIHOX4*, *PAX3/7*, neurotrophic tyrosine kinase receptor precursor (*NTKR*), *NOTCH1L* and *NOTCH2*. Notably, OSMs regulated putative target genes encoding proteins involved invertebrate brain function and nerve disease, visual sensing in the nerve cord, including adult brain protein 239-like, huntingtin and opsin protein.

In the muscle, OSM predicted targets related to ubiquitination are overrepresented, including E3 ubiquitin-protein ligase family (*E3 ubiquitin-protein ligase HUWE1-like*, *E3 ubiquitin-protein ligase CBL-like*, *E3 ubiquitin-protein ligase MARCH6* and *E3 ubiquitin-protein ligase MURF2*) and ubiquitin conjugation factor E4 B (*UBE4B*). In the intestine, hepatic caecum and gill, a large number of genes involved in the innate immune system are regulated by OSMs, including HSPs, complement component, PRRs, cytokines, apoptosis-related proteins, adaptors, signal transducers and caspases; particularly, mucin-2 (*MUC2*), a major mucin of the colon mucus in the vertebrates, frequently appeared to be potentially regulated targets of OSMs in the intestine and hepatic caecum. Interestingly, in the list of predicted target genes of OSMs in the hepatic caecum, we noted some insulin-related genes, such as insulin-degrading enzyme (*IDE*), insulin-like growth factor and insulin-like growth factor 1 receptor (*IGF1R*).

### Validation of miRNA expression profiles using qRT-PCR

To further validate the expression of the identified miRNAs by deep sequencing, the expression patterns of eight randomly selected known miRNAs across different organs, were investigated using Taqman miRNA probes. Linear correlation analyses of the fold-changes in the relative expression levels between the miRNA sequencing and quantitative real-time PCR (qRT-PCR) results showed a significant correlation for each miRNA (Pearson’s correlation coefficients > 0.8, *P* < 0.05) (Figure 3D).

## DISCUSSION

In this study, approximately 34 million clean reads across eight organ of *B. belcheri* were generated, which is the largest so far obtained for sRNAs of *B. belcheri*, allowing us to obtain adequate miRNA molecules that are contained in different organs. Additionally, the miRNA expression levels quantified by qRT-PCR analysis of eight randomly selected known miRNAs indicated that deep sequencing is reliable, which provided a quality guarantee for downstream bioinformatics analysis. A total of 190 miRNAs (167 known) were detected in the current study. Zhou *et al*. (2017) performed systematic investigation of *B. floridae* microRNAs using a computational pipeline to predict miRNAs from throughout the genome. The authors identified 245 predicted miRNAs, which is more than the present study. The higher number of predicted miRNAs can be attributed to false positives resulting from only using a bioinformatic genome scan under a regular threshold. More importantly, miRNAs identified in the current study can indeed be expressed in organs of adult amphioxus. However, a proportion of those detected by genome-wide bioinformatic scan may not be expressed or only expressed at low levels in adult animals under normal conditions. Some examples could be those miRNAs involved in development, immunology, and response to stress. In addition, a higher proportion (79/190 = 41.58%) of miRNAs were predicted to be expressed in an organ-specific manner. By contrast, a previously published study reported that only 47/245 (19.18%) were development-specific miRNAs (Zhou *et al*., 2017). This could be the result of different algorithms being used to detect specific miRNAs. Alternatively, organ function formation depend on more specific miRNA than developmental processes in amphioxus.

The vast majority of the conserved miRNAs detected here have already known been annotated (only novel_mir1 was a newly detected miRNA). Previous studies showed that an average transcripts per million (TPM used for quantification of expression level) values of the top 30 known miRNA identified in *B. belcheri* averaged 2272.4. By contrast, TPM values of the top 30 novel miRNAs was only is 131.1 (Zhang *et al*., 2017a). The conserved miRNAs present higher expression levels than *B. belcheri*-specific miRNAs, and they were detected as known miRNAs more easily, because of higher abundance of known miRNAs than that of novel miRNAs in amphixous. Furthermore, the majority of the OSMs detected here were found to be conserved with *B. floridae* and other vertebrates. This suggests that amphioxus OSMs play a key role in organ function formation in amphioxus speciation, and contributing to the complexity of the vertebrate body. Interestingly, conserved functions of the miR-1/miR-133 cluster in chordate muscle differentiation have been proposed, and their specific location was preliminarily assessed by *in situ* hybridization using amphioxus embryos (early, early-mid, and late neurulae) and early larvae (Campo-Paysaa *et al*., 2011). In the current study, muscle-specific expression of all miR-1 and miR-133 OSMs (e.g. bbe-miR-133-5p, bbe-miR-133-3p and bbe-miR-1-5p) was observed. These observations lend additional support to the hypothesized role of miR-1 and miR-133 in amphioxus muscle differentiation. Moreover, i*n situ* hybridization of miR-1 and miR-133 produced spatial expression data similar to that obtained through miRNA sequencing. This demonstrated the reliability of the analyses conducted in this study. The conserved novel_mir1 identified in this study is homologous to bfl-miR-4871-3p. However, the function of the miR-4871 family remains elusive. Analysis of predicted target genes showed that this conserved OSM regulates genes encoding fibroblast growth factor receptors (FGFRs) that promote the proliferation and differentiation of various cells via binding cell surface-expressed receptor tyrosine kinases (Han *et al*., 2017). Therefore, the novel_mir1 may contribute to skin function formation in amphioxus, although further research is needed for confirmation.

The data presented here suggests show that some OSM target genes enriched in certain GO terms are limited to organ-specific functions. Those involving epidermis development and regulation of epithelial cell proliferation were detected in the skin. In the ovary, GO terms with organ-specific functions are related to gonad development. Muscle -specific GO terms in the muscle are primarily associated with muscle cell and organ development, which is consistent with data obtained using porcine and human skeletal muscle (Baccouche et al., 2014; Martini *et al*., 2014). Taken together, the data suggests that OSM regulatory functions in amphioxus muscle are conserved in vertebrates. For those OSMs in the intestine, their predicted target genes are also enriched to organ-specific GO terms related to the proteolysis. Proteolysis is the breakdown of proteins into shorter peptides and amino acids to aid in the digestion of food in the intestine (Aloğlu *et al*., 2011). Therefore, proteolysis would be expected to be a key intestine-specific function among various amphioxus organs. It was also observed that some hepatic caecum -specific GO terms, such as those involving organic metabolism are functions that have also been associated with the liver in other vertebrate species (Araújo *et al*., 2018). A large number of nerve cord-specific GO terms involving nervous system-related functions and development were represented, indicating that the nervous system of amphioxus is unexpectedly functionally complex with diverse function, as previously described (Benito-Gutiérrez, 2006). Interestingly, some biological processes managed by the vertebrate brain are exhibited specifically, such as circadian rhythm, learning, sleep, light response and vision. Previous publications have proposed that the anterior of the amphioxus cerebral vesicle (neurula) is homologous to the thalamus, pretectum (as part of diencephalon that is part of the forebrain in vertebrates) and midbrain of vertebrates based on the expression pattern of developmental protein-coding genes (such *HOX*, *OTX*, *DLX*, etc.) (Holland and Holland, 1998; Albuixech-Crespo *et al*., 2017). Here, the analysis supports findings that the vertebrate brain originates from part of the ancestral nerve cord based on the perspective of miRNA expression. In the gill and notochord, GO terms involving a wide range of biological functions are overrepresented. Particularly, innate immune-related GO terms, such as those involving cell apoptosis, immune system, and defense response were widely distributed, especially in the intestine, gill, and hepatic caecum. This wide organ distribution of immune-related GO terms may be explained by the fact that these tissues are on constant contact with seawater, which is the source of most microbial exposures.

The bbe-miR-182 family was detected as known OSMs in the amphioxus skin. As reported in previous studies, miR-182 expressed at specific levels in the skin is associated with breast cancer and basal cell carcinoma (Sand *et al*., 2012; Wang *et al*., 2013), suggesting that miR-182 is an ancient miRNA family of skin-specific regulators. Furthermore, it was predicted that a miR-182 may be a key regulator of innate immune- and calmodulin-related genes, thus miR-182 may be a potential molecular indicator of immune system activities in the research of skin immunity of amphioxus. At the time of publication, no reports on the function of miR-2068 and miR-4873 families could be found. Here, it was observed that, in the ovary, it was predicted that the OSMs miR-2068 and miR-4873 regulate *PIWIL1*, which belongs to the piRNA pathway (Rengaraj *et al*., 2014). These observations suggest potential regulatory function of miRNAs to piRNA and the ovary immune system in amphioxus. Despite notochord-specific GO terms are not being found, some genes involved in notochord development were predicted in the target gene analysis. These OSMs may be candidates for investigation of notochord evo-devo. It was observed that not only known OSMs that potentially target genes involving vertebrate brain function, as well as development and diseases of the nerve system were predicted, but also those that regulate typically innate immune-related genes. For example, Liu *et al*. (2016) demonstrated that amphioxus LGP2, a target of bbe-miR-4928-3p as OSMs, plays an antiviral role similar to that observed in other vertebrate species by comparing expression in *Branchiostoma japonicum* challenged with viral mimic poly(I:C) and untreated controls. These results provide evidence that neuro immunity is active and that miRNAs may be mediators of the neuro-immune system in early branching chordates. In the review by Candiani *et al*. (2012), the role of mir9, mir124, mir219 in deuterostomes were possibly associated with evolution of a central nervous system (CNS). In support of the above review, nerve cord-specific expression of these three miRNA families were observed in this study. Target genes of mir9, mir124, and mir219 were predicted using bioinformatics. In addition, several known and novel miRNAs with nerve cord-specific expression were newly identified in the current investigations. These data serve as a foundational genetic resource to support future exploration of CNS evolution in chordates. In the muscle, known OSMs (e.g. bbe-miR-1-5p, bbe-miR-133-3p, bbe-miR-2058-5p and bbe-miR-4859-3p) are primarily involved in ubiquitination. The E3 ubiquitin ligases could function in ubiquitin-mediated muscle protein turnover, promoting skeletal muscle differentiation and myofibrillogenesis (Perera *et al*., 2012). Furthermore, miR-133 and miR-1 were found to be muscle-specific miRNAs conserved in vertebrates (Tani *et al*., 2013). The data presented here show muscle-specific expression of these two miRNA families, revealing their conservation in chordates, not just in higher vertebrates. Many studies have discussed the key roles of the intestine, gill, and hepatic caecum as the most important immune organs in the defense against various insults in amphioxus, including miRNA, organ structure, and protein-coding genes (Han *et al*., 2010). Here, OSMs involved in immune function in these three organs were identified, suggesting that the immune system may be active and regulated by miRNAs in multiple organss of amphioxus. Notably, insulin-related enzyme and growth factor (*IDE*, *IGF1*, *IGF1R*), as predicted targets of OSMs in the hepatic caecum, are well known for playing specific key roles in the pancreatic cells of vertebrates (Zhang *et al*., 2007; Fernández-Díaz *et al*., 2018). Therefore, in later branching chordates, it is reasonable to speculate that the ancestral homologous organs of vertebrate pancreas probably have functional connection to a certain extent with the hepatic caecum.

In summary, the identification and quantification of miRNA in different organs is a first key step in the investigation of their associated functions in amphioxus. Deep sequencing was employed to detect and quantify miRNAs in eight organs. Bioinformatic analysis was used to detect OSMs based on these miRNAs. Many of the miRNAs identified here had not been previously reported. Of note, due to a largely unknown and complex miRNA functions, the regulatory roles of several OSMs in this study could not be concluded. This is the first study to explore the dynamics of miRNAs in multiple organs at a genome-wide level using deep sequencing in amphioxus. Our analysis contributes to the field’s understanding of the differentiation of organ function in amphioxus.

## COMPLETING INTERESTS

The authors declare no competing financial interests.

## ACKNOWLEDGMENTS

This study was supported by the foundation of Guangxi Key Laboratory of Beibu Gulf Marine Biodiversity Conservation, Beibu Gulf University (No: 2019KA01 and 2019ZB09), and the Scientific Research Foundation Project of Yunnan Education Department (2019J0050), and Natural Science Foundation of Guangxi Province (No: 2016GXNSFBA380156, 2016GXNSFCA380007), and the National Natural Science Foundation of China (No: 31760713).

## SUPPLEMENTARY MATERIAL

**Supplementary File 1.** Taqman probe information of randomly selected miRNAs used in qRT-PCR analysis.

**Supplementary File 2.** Summary of tags generated from sequencing of eight organs small-RNA libraries in *Branchiostoma belcheri*.

**Supplementary File 3.** Length distribution of adult *Branchiostoma belcheri* small RNA (sRNA) in each of eight organs. X axis is length andY axis is number of clean reads. Warmer color means shorter length of sRNA, while cooler color refer to longer. **Supplementary File 4.** The stacking histogram for annotation of small RNAs in each of eight organs. Exon and intron < 3% tags generated by the degradation of mRNAs. **Supplementary File 5.** MicroRNAs homologous with those of vertebrates and *Branchiostoma floridae* in adult *Branchiostoma belcheri*.

**Supplementary File 6.** List of novel miRNA and their precursors identified newly in eight organs of adult *Branchiostoma belcheri*.

**Supplementary File 7.** Predicted precursor structure of all the novel miRNAs identified in this study.

**Supplementary File 8.** List of organ-specific miRNAs identified in each organ of adult *Branchiostoma belcheri*.

**Supplementary File 9.** List of the GO enrichment terms of targeted genes of organ-specific miRNAs in each organ of adult *Branchiostoma belcheri*.

**Supplementary File 10.** The predicted targeted genes of organ-specific expressed known miRNAs in each organ of adult *Branchiostoma belcheri*.

